# Crystallographic fragment screen of Enterovirus D68 3C protease and iterative design of lead-like compounds using structure-guided expansions

**DOI:** 10.1101/2024.04.29.591650

**Authors:** Ryan M. Lithgo, Charles W.E. Tomlinson, Michael Fairhead, Max Winokan, Warren Thompson, Conor Wild, Jasmin Cara Aschenbrenner, Blake H. Balcomb, Peter G. Marples, Anu V Chandran, Mathew Golding, Lizbe Koekemoer, Eleanor P. Williams, SiYl Wang, Xiaomin Ni, Elizabeth MacLean, Charline Giroud, Tryfon Zarganes-Tzitzikas, Andre Schutzer Godoy, Mary-Ann Xavier, Martin Walsh, Daren Fearon, Frank von Delft

## Abstract

The development of effective broad-spectrum antivirals forms an important part of preparing for future pandemics. One cause for concern is the currently emerging pathogen *Enterovirus D68* (EV-D68) which primarily spreads through respiratory routes causing mostly mild to severe respiratory illness but, in severe cases, acute flaccid myelitis. The 3C protease of EV-D68 (3C^pro^) is a potential target for the development of antiviral drugs due to its essential role in the viral life cycle and high sequence conservation amongst family members. In this study, we describe the identification of fragments which bind to3C^pro^ using crystallographic screening and the expansion of these into more lead-like compounds. The hits revealed interesting directions for hit-to-lead progression, specifically the importance of the pocket occupied by the conserved glutamine sidechain of the substrates and the interactions formed. Additionally, two pockets could be joined by not following the backbone of the native substrates, thus circumventing the screening issues arising from the flexibility of the catalytic triad. These observations of the novel binding modes of the chemical matter found by this screen can help shape future drug design campaigns against 3C proteases.

## Introduction

The development of novel, low cost and globally available antiviral therapeutics remains an essential goal to ensure we are prepared for future pandemics. Although researchers were able to develop and deploy safe and effective vaccines at an unprecedented speed, more than 6.9 million lives were lost to SARS-CoV-2 [50]. This mortality rate would have been vastly reduced had easily deployable, broad spectrum oral antivirals targeting coronaviruses been available. Such broad-spectrum antivirals are an important strategic part of combating viral diseases and reducing morbidity and mortality [3, 38] and the timely availability of effective antivirals could, therefore, save millions of lives in the next pandemic.

Enterovirus D68 (EV-D68), a *Picornavirus*, is an emergent pathogen that primarily infects children causing mild to severe respiratory illness and in some instances acute flaccid myelitis, a serious neurological condition[14]. EV-D68 is known to spread primarily through respiratory routes as well as faecal-oral routes and is, hence, a concern for pandemic spread if infectivity were to increase [1]. In recent years, large outbreaks have been reported worldwide with a case in the United States in 2014 causing severe illness for more than one thousand patients and likely many more undiagnosed mild or asymptomatic infections [37]. Antiviral research efforts have been made to develop EV-D68 inhibitors targeting the viral life cycle such as replication, translation and polyprotein processing [20]. However, there are currently no vaccines or antivirals available to prevent or treat EV-D68 infection.

During the replication of EV-D68, the viral genome is translated into a polyprotein which is processed by the 2A, 3C and 3CD proteases. EV-D68 3C protease (3C^pro^) is a cysteine protease that has two domains in a chymotrypsin-like fold with a Cys-His-Glu catalytic triad in the active site [45]. The function of 3C^pro^ cysteine protease is to process the viral polyprotein through the cleavage of the non-structural proteins (NSPs) at their N-termini. These substrate sequences contain several highly conserved residues which are key to 3C^pro^ activity (Figure 1). 3C^PRO^ additionally targets multiple host proteins that ultimately promote infection and viral replication, whilst simultaneously suppressing host immune responses [44, 54, 55].

**Figure 1:**
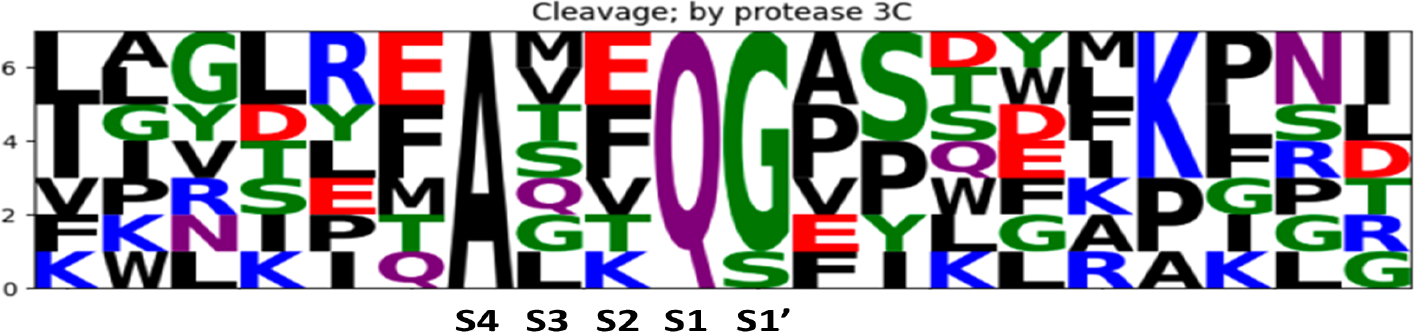
Sequence logo of the 3C^pro^ substratesequences found within the viral polyprotein, with the position they adopt in the active site shown. The cleavage of the substrate occurs between the S1 and S1’ sites, whereby the S1 amino acid is universally glutamate (Q) and S1’ is either a glycine or serine. The S2 and S3 amino acids vary between the different substrates, however the S4 site must contain an alanine.

The overall protein sequence for 3C^pro^ is heavily conserved across multiple species of picornavirus’, with the residues that make up the entire catalytic site conserved the most (See **Fig.2**). The high levels of sequence conservation leads to the possibility that broad-spectrum pan-picornovirus 3C^pro^ inhibitors can be developed.

**Figure 2:**
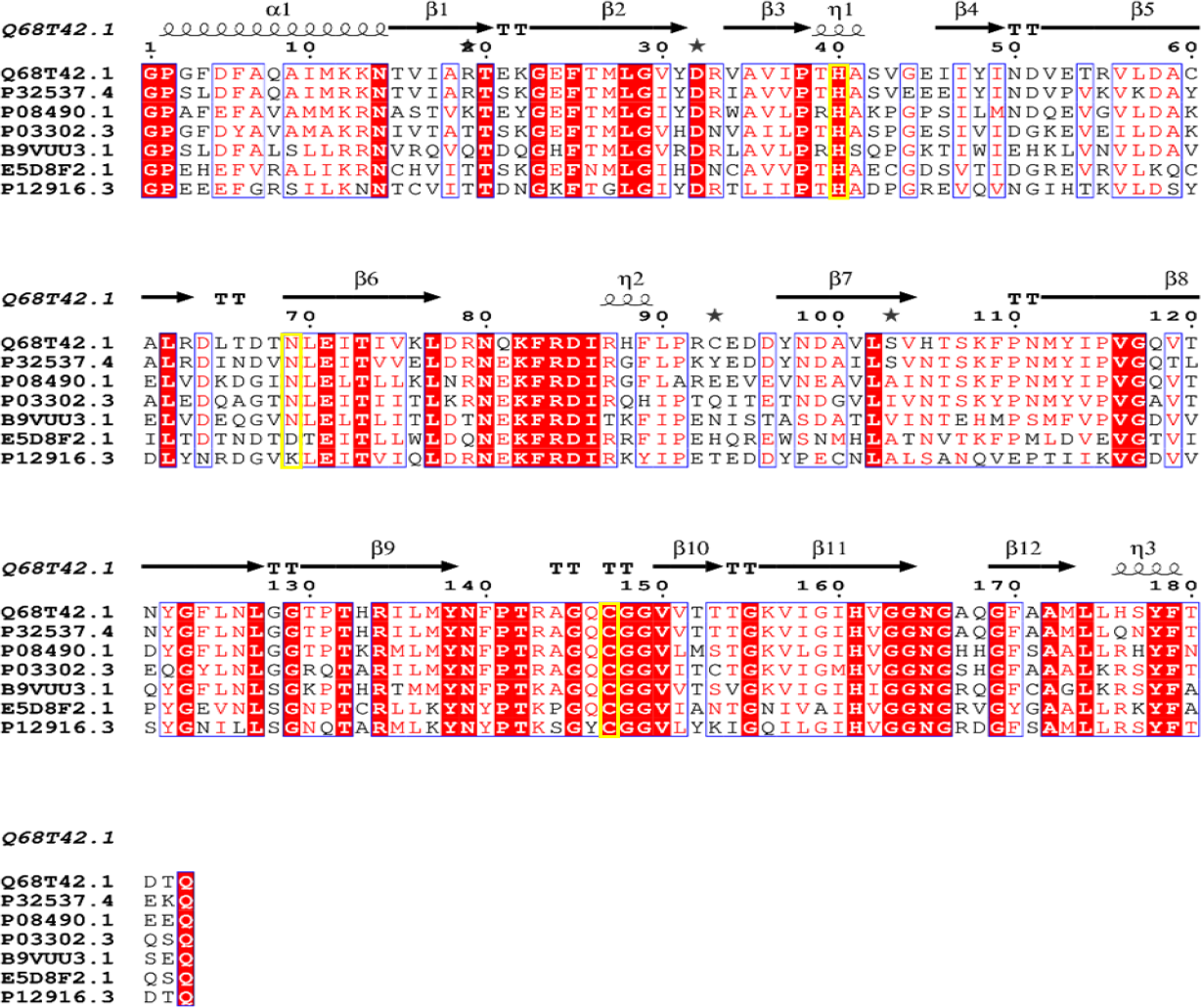
*Sequence alignment of several picornovirus* 3C^pro^ sequences; Human Enterovirus D68 (EVD68-3C^pro^) (Q68T42.1), D70 3C^pro^ (P32537.4), Human Enterovirus 71 3C^pro^ (B9VUU.3), Echo 9 virus 3C^pro^ (P08490.1), Poliovirus type 3 3C^pro^ (P03302.3), Human Rhinovirus C 3C^pro^ (E5D8F2.1), 1B 3C^pro^ (P12916.3). Conserved residues are highlighted in red, with the catalytic triad residues highlighted by a Yellow box. Regions of the greatest conservation surround the active site residues.

Fragment-based drug discovery (FBDD) is a proven method for generating hits as starting points for drug discovery. It differs from high-throughput screening (HTS) by analysing smaller libraries (500-2,000 vs 100,000 compounds) with simpler molecules(fragments vs lead-like compounds) which have on average a smaller molecular weight (<250 Da vs >300 Da) [27, 40]. Sensitive detection methods are needed to reveal the weak but highly efficient binding fragments.s. Crystallographic fragment screening is one such highly sensitive method that relies on direct detection of bound fragments within electron density data [22]. The obtained structural information can immediately be used to guide further optimisation of fragment hits, in a rapid design-make-test cycle. Popular strategies for fragment elaboration include fragment growing, fragment linking, and fragment merging which can quickly result in potent lead-like compounds. [10]. In the case of FBDD via crystallographic screening, the compounds sharing similar bound-conformations, interactions and binding modes can be leveraged by identifying derivative compounds that preserve these. The design of these compounds can be traditional medicinal chemistry, or we can leverage the increasing effectiveness of algorithmic design.

Here we present the results of a crystallographic fragment screen against 3C^pro^. In total, 1,231 fragments were screened resulting in 101 bound compounds across 3 main sites, including the catalytic site. These hits were used as starting points to design new novel follow-up compounds in areas that have been unexplored previously and move away from the literature based peptidomimetic inhibitors. The follow-up compounds were designed using multiple iterative cycles of various algorithmic designs, utilising fragment merging, linking and growing approaches that recapitulated the parent fragment hit interactions. The compounds designed using these algorithms were curated and selectedfor synthesis by expert medicinal chemists before being analysed using high-throughput crystal soaking, with the results feeding back into the iterative process of further designs.

## Materials and methods

### 3C^pro^ construct design

A synthetic gene, codon optimised for expression from *E*.*coli*, corresponding to residues 1549-1731 of the Enterovirus D68 polyprotein (Uniprot id: Q68T42.1) was ordered from TWIST Bioscience. TThe gene was cloned using the Golden Gate method [15] into a pNIC vector (https://www.addgene.org/215810/) with a non-cleavable C-terminal hexaHIS tag (bold - below). The final construct sequence is:

MGPGFDFAQAIMKKNTVIARTEKGEFTMLGVYDRVAVIPTHASVGEIIYINDVETRVLDACALRDL TDTNLEITIVKLDRNQKFRDIRHFLPRCEDDYNDAVLSVHTSKFPNMYIPVGQVTNYGFLNLGGTP THRILMYNFPTRAGQCGGVVTTTGKVIGIHVGGNGAQGFAAMLLHSYFTDTQK**HHHHHH**

Plasmid is available at Addgene: https://www.addgene.org/204817/

### Expression and purification of 3C^pro^

Expression of 3C^pro^ consisted of transforming the plasmid into *E. coli* BL21(DE3)-RR, and cells were grown at 37°C in TB medium supplemented with kanamycin (50 μg/mL). After reaching an optical density at 600 nm of around 1.8, the temperature was lowered to 18°C before induction of protein expression overnight by adding 0.5 mM IPTG. Harvested cells were resuspended in lysis buffer [10 mM HEPES (pH 7.5), 500 mM NaCl, 5% glycerol, 30 mM imidazole, 0.5 mM TCEP, 1% TX-100, 0.5 mg/mL lysozyme, 0.05 mg/mL benzonase]. Proteins were first purified by immobilised metal affinity chromatography (IMAC) using Ni-Sepharose 6 FF resin (Cytiva). The column was washed with binding buffer [10 mM HEPES (pH 7.5), 500 mM NaCl, 5% glycerol, 30 mM imidazole, 0.5 mM TCEP] and target protein eluted using same buffer containing 500 mM imidazole. Protein was lastly purified by Size-Exclusion Chromatography (SEC) (SRT SEC-100, Sepax) in SEC buffer (10 mM HEPES (pH 7.5), 500 mM NaCl, 5% glycerol and 0.5 mM TCEP). Proteins were characterised by SDS-polyacrylamide gel electrophoresis and mass spectrometry before flash-freezing in liquid nitrogen for storage at -80°C. The full protocol can be found here https://dx.doi.org/10.17504/protocols.io.dm6gpzrkdlzp/v1

### 3C^pro^ Fluorescence Resonance Energy Transfer (FRET) Assay

The proteolysis activity of purified 3C^pro^ was tested by performing IC_50_ measurement of the control inhibitor GC376 (Pubchem CID 71481119) in a biochemical FRET assay. GC376 was dispensed onto 384-well, small volume black plate (Greiner, #CLS3544) using the Echo liquid dispenser (Beckman

Coulter) at a final top concentration of 50 µM using a dilution factor of 1.5. Ten microliters of 2x (1 uM) EV-D68 3C solution (50 mM Tris pH 7.0, 150 mM NaCl, 10% glycerol and 1 mM TCEP) was added to dispensed GC376 and incubated for 1 hour at room temperature. The reaction was initiated by the addition of 10 µL of 2x (40 µM) substrate solution (Dabcyl-KEALFQGPPQFE-Edans (LifeTein, USA)) made in reaction buffer. The fluorescence intensity at 460 nm was read every 30 seconds for 2 hours in kinetic mode, which included a shaking step of the plate between each measurement. The IC_50_ was calculated by plotting the initial velocity against various concentrations of the tested inhibitor by using a four parameter dose−response curve in Prism (v8.0) software. The Z’ or Z-factor was calculated as below (μ: mean; σ: standard deviation, s: no inhibitor signal, highest inhibition signal). Full protocol: https://dx.doi.org/10.17504/protocols.io.261ge54jyg47/v1

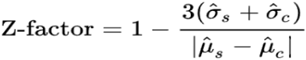

### Crystallisation of 3C^pro^

Purified 3C^pro^ in SEC buffer was screened in Shotgun (SG1) (Molecular Dimensions) and Index (Hampton Research) screens at a concentration of 1 mM (35 mg/mL). Screens were set up in SwissSci 3 drop plates, with drops set up at 3 ratios 1:2, 1:1 & 2:1 (protein:reservoir) to a total volume of 300 nl. Once prepared screens were incubated at 20°C with images taken after 24 hrs and then following a fibonacci sequence of timings.

Crystals of 3C^pro^ grew within the Index screen in condition G9 (0.2 M Ammonium acetate, 0.1 M Tris (pH 8.5) & 25% w/v PEG 3350), at a ratio of 1:2 (protein:reservoir) after approximately 2 days. Crystals were mounted without cryoprotectant using 150 um loops and flash frozen in liquid nitrogen. Data collection was conducted on I04-1 at Diamond Light Source (UK) at 100K, with the data processed using Autoproc [46]. The structure of 3C^pro^ protease was processed to 1.5 Å with a spacegroup of P 2_1_ 2_1_ 2_1_, with the structure deposited on the PDB (8CNX). A table of data collection and refinement statistics can be found in Supplementary 1.

Optimisation of the crystal system was carried out by creating a screen around the original crystallisation condition. The optimisation screen varied the pH (ranging between 7.5 - 9.0) and the PEG 3350 percentage (ranging between 18 - 32 % w/v), crystals grew in most conditions after approximately 2 days with the best diffracting data came from the condition 0.2 M Ammonium acetate, 0.1 M Tris (pH 8.14) & 25% PEG 3350 w/v, with a protein:reservoir ratio of 100:200 nL and was determined to be the condition to use going forward.

To maximise crystal reproducibility, crystal seeding was also implemented. Using D68 3C crystals from the optimisation screen a solution several seed dilutions were prepared following the protocol outlined here (https://dx.doi.org/10.17504/protocols.io.kxygx3nwog8j/v1). 50 nL of crystal seed was added to the 300 nL crystallisation drop, with crystals growing in all conditions. The seed stock that gave reproducible crystals which were large in size and diffracted readily (∼1.5 Å) grew using a 50 nL addition of 1:100,000 seed dilution.

The full crystallisation protocol can be found at: dx.doi.org/10.17504/protocols.io.5qpvoky29l4o/v1

### Crystallographic fragment screen of 3C^pro^

Fragment soaking was conducted using acoustic dispensing from an Echo 650 liquid handler (Beckman, USA). Dispensing of the fragments was targeted for regions of the crystal drop that were as far from the crystals as possible, this reduced any potential liquid transfer induced damage [8]. Prior to soaking with fragments the DMSO tolerance of the crystals was determined by varying DMSO concentrations between 0-30 % at 5 % increments, and the time of soaking between 1 and 3 hours. After soaking crystals were mounted using a Oxford Lab Technologies crystal shifter [47] and X-ray diffraction data collected on I04-1 (DLS). Based on the diffraction data the crystals were deemed to tolerate 25% DMSO for 3 hours.

Fragment screening consisted of soaking 1,231 fragments from several libraries from the XChem facility at Diamond Light Source including the DSi-poised, DSi-poised EUbOpen [9 & 16], MiniFrags [36], Fraglites [40], the PepLite library [11], York3D [12] SpotXplorer [5] and Covalent MiniFrags library [24, 25, 26]. Fragment solutions (50 nL, 12.5% of final drop volume) were dispensed into the 300 nL crystallisation drop using the Echo, plates were resealed and incubated at 20°C for 2.5 hours after which the crystals were mounted and stored in liquid nitrogen without the addition of a cryoprotectant.

Data collection was carried out using I04-1 at Diamond Light Source (UK) at 100K, with the data automatically processed using XDS [23], Autoproc [46], Xia2 [50] or DIALS [51]. The majority of D68 3C^pro^ structures were processed to a high resolution of between 1.5 and 1.7 Å. Further downstream analysis was carried out using the XChemExplorer pipeline [28], with electron density maps generated using DIMPLE [45] and ligand binding events identified using PanDDA [39] and PanDDA2 [48]. Ligand restraints were generated using grade [51], with ligands modelled into the PanDDA/PanDDA2-calculated event maps using Coot [13]. All structures were refined using Refmac [35]. Data collection and refinement statistics are summarised in data file S1.

Coordinates, structure factors, and PanDDA event maps for the structures discussed are deposited in the PDB (group deposition G_1002271) and structural data is also available through the Fragalysis webserver.

### 3C^pro^ Fragment Elaborations

#### Fragmenstein

[31] is operated in two modalities: combination or placement. In the combination route atoms are merged by their coordinates and the stitched together compound corrected and energy minimised. In the placement route a compound is mapped against one or more hits via several rules and an interactive maximum common substructure approach.

#### FragmentKnitwork

enumerates compounds that incorporate substructures from two parent hits [49] or shape-and-colour analogues using a graph database search of catalogue compounds. These compounds were placed with Fragmenstein.

#### Arthorian Quest

[https://github.com/matteoferla/Arthorian-Quest] addresses specific design ideas. Ambiguous SMARTS patterns from parent compounds are generated via human choice. Their dot-separated concatenation is used to search in a NextMoveSoftware Arthor server hosted by Prof John Irwin (https://arthor.docking.org/) [21]. These derivatives are placed with Fragmenstein against the parent hits. The design ideas were expansions of terminal S1 pocket hits towards the catalytic cysteine (including the displacement of oxyanion water) and to bridge S1 and S2 hits via a variety of hypotheses.

#### OpenEye ROCS

carries out shape-and-colour based searches. A query was manually built for the P2 pocket as this is a hydrophobic pocket.

#### STRIFE

is a deep learning tool for *de novo* compound generation which leverages identified binding hotspots for its 3D aware expansions.

The compounds generated by Fragemstein (combination route) and STRIFE were used as queries in an analogue search using a NextMoveSoftware SmallWorld server hosted by Prof John Irwin (https://sw.docking.org/) [20]. These were then placed with Fragmenstein against either the parent hits or the design.

The enumerated compounds, in most instances, were ranked by a multifactor metric taking into account X, Y, Z. These were then k-mean clustered by residue-level interaction presence scaled by frequency of that interaction in the whole dataset.

#### Fragalysis - data curation and exploration tools

the top compounds were visually assessed prior to purchase from Enamine Ltd.

## Results and discussion

### 3C^pro^ FRET Assay

The catalytic activity of purified 3C^pro^ was confirmed by FRET assay by measuring the IC_50_ of GC376, a viral 3C^pro^ inhibitor already characterised in the literature [30]. The initial velocity measured in the linear range of the reaction was normalised and plotted against GC376 concentration. The assay shows a robust signal characterised by a signal-to noise of 937 and a Z’ of 0.795 and allowed a consistent and expected IC_50_ of 0.26 µM (Figure 3). The results of the assay show that the 3C^pro^ construct designed and purified is active, giving confidence that what is observed in the crystallographic structures of the fragment screen are biologically relevant.

**Figure 3:**
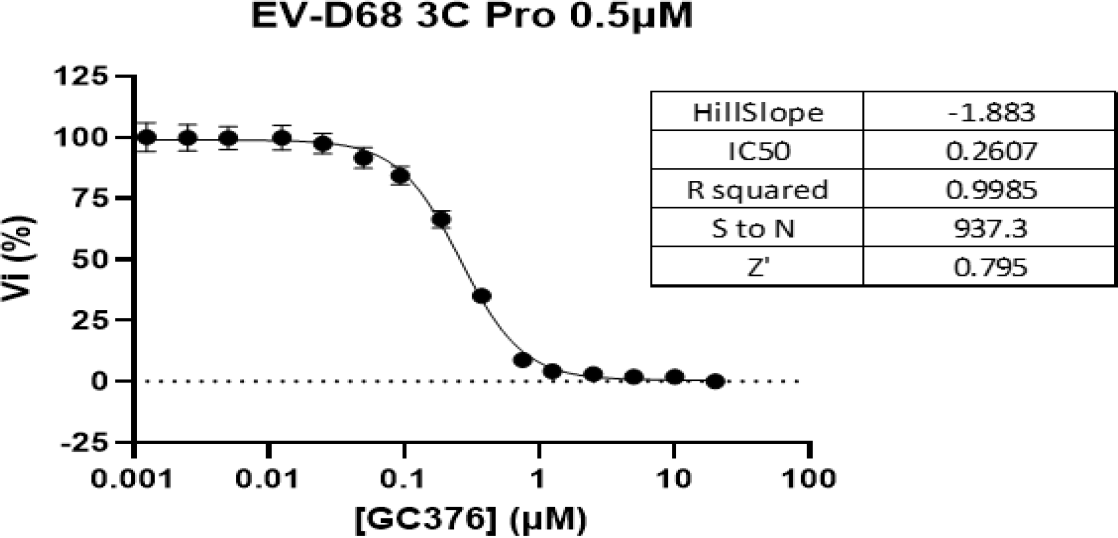
Protease assay of 3C^pro^ against literature inhibitor GC376. The assay showed a IC_50_ of 0.26 µM which is expected for the compound.

### Structure of apo 3C^pro^

We solved the apo crystal structure of 3C^pro^ at 1.5 Å, showing two copies (monomers A & B) within the asymmetric unit (ASU) (PDB ID: 8CNX). The structure displays the typical chymotrypsin-like fold consisting of two β-barrel domains typically found in serine proteases [12]. Both domains form a long shallow groove for substrate binding, with the Cys-His-Glu catalytic triad active site, similar to the Ser-His-Glu of serine proteases [34], found at the interface of both domains (See Figure 4). The active site is composed of 5 sub-units (S) in a ‘+’ shape; S1’, S1, S2, S3 & S4 [29]. The S1 site of 3C^pro^ forms a polar channel within which the highly conserved glutamine of the substrate sits. His161 which is positioned directly under the catalytic Cys147, is ideally positioned to hydrogen bond the carbonyl of glutamate within the substrate. Directly adjacent to this is Thr142 which contains a polar hydroxyl group which can hydrogen bond to the amine head of the glutamine substrate. In addition Thr142 is involved in a heavily coordinated water molecule, also coordinated by the carbonyl backbones of Phe140 and Gln168. This water may be involved in the specificity and binding of the glutamine of the substrate. The S1’ region of 3C^pro^ contains the oxyanion hole brought about through the highly conserved Gly-X-Cys-Gly-Gly motif within the active site, with the NH groups of both Gly145 and Cys147 forming the hole. The hole stabilises the transition state during proteolytic cleavage [34]. The structure we obtained showed the presence of the hole through the positioning of Gly145 and Cys147 with a water molecule heavily coordinated in the structure [45].

**Figure 4:**
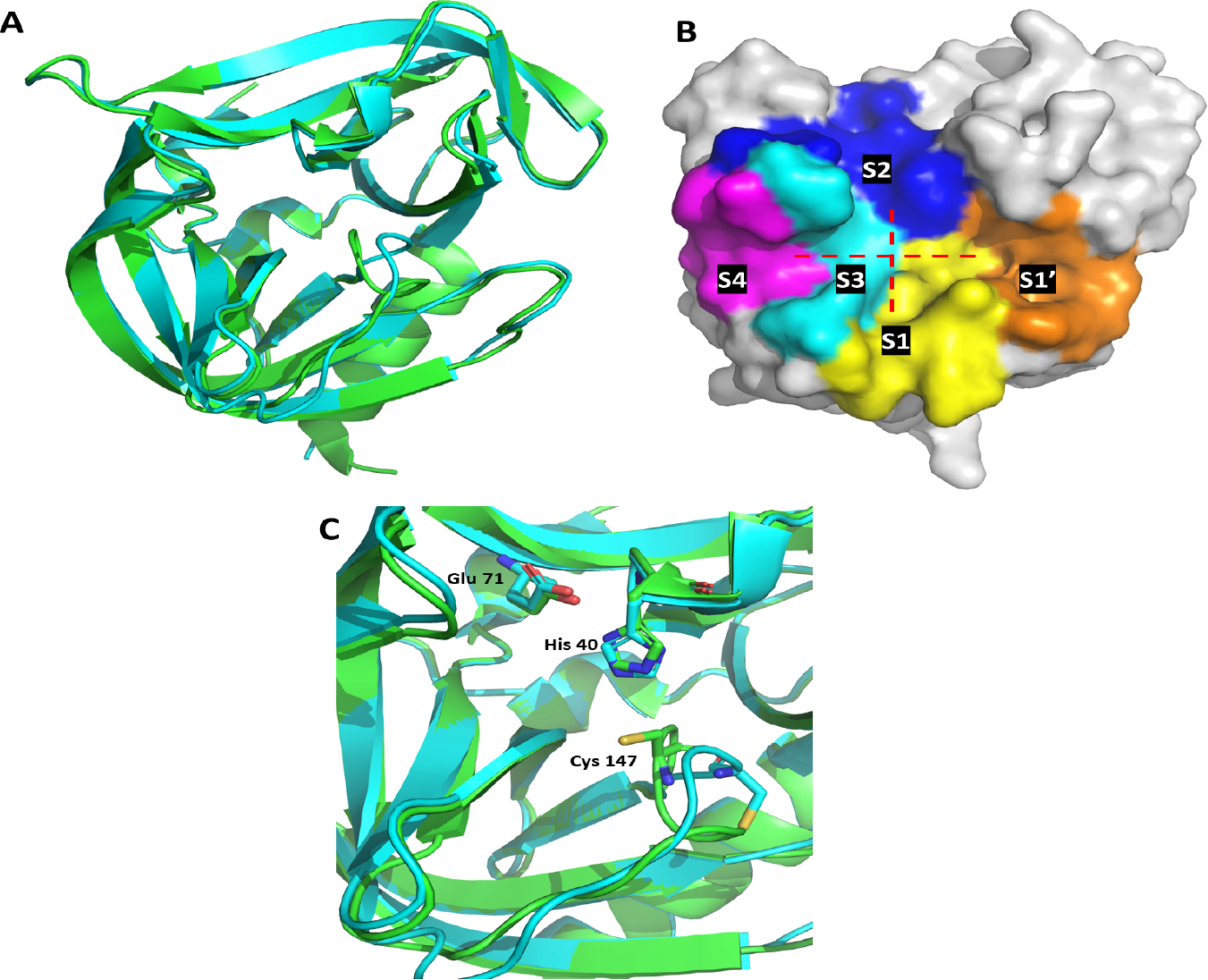
**(A)** Overlaid cartoon illustration of the crystallographic structure of 3C^pro^, showing monomers A (green) and B (cyan) with their secondary structures shown. **(B)** Surface representation of 3C^pro^ monomer A highlighting the active site subunits in various colours with the ‘+ like’ shape of the active site highlighted in dashed red lines. The S1’ (orange) site is positioned on the right hand side of the active site where the conserved glycine/serine is located. The S1 (yellow) site is positioned at the bottom of the active site and forms a cleft where the substrate glutamine would be positioned. The large S2 (blue) site is at the top of the active site. The S3 (cyan) and S4 (pink) sites are to the left of the active site and is where the conserved alanine resides (S4). **(C)** The active sites of monomer A & B, showing the similarities in the positions of His40 and Glu71. The positioning of Cys147 in monomer A can be seen to adopt the atypical Ser-His-Glu triad formation, however monomer B shows that Cys147 is everted 180° away from the active site and results in the entire loop adopting a different conformation.

The S2 region of the active site within 3C^pro^ is large and would accommodate the variety of substrate amino acids that can bind there, the position of the His41 may enable some form of pi-stacking to occur. The position of His40, adjacent to Cys147, acts as the acid-base catalyst for the Cys-His catalytic diad. The negatively charged carboxylate of Glu71 within the S2 region is generally thought to be required for stabilising the productive conformation of the histidine and the active site architecture, this is observed in the structure that we obtained [6, 10, 12].

The S3 region is rather constrained and supports previous observations that the amino acid sidechain of the substrates tend to be orientated into the solvent exposed region. Finally the S4 region of the active site is considerably hydrophobic, consisting of Leu125, Ile135 and Phe170, the hydrophobicity of this region compliments the alanine that is highly conserved within the substrate sequences.

Although monomer A conforms to the anticipated Ser-His-Glu active site conformation, the active site of monomer B does not. We observe that the Cys147 sidechain is everted 180° away from the active site disrupting the catalytic triad. The observed differences between the two monomers in the apo structure is not limited to Cys147, the entire loop that positions the cysteine, ranging from residues Asn139 to Gly149 adopts a completely different conformation in each monomer. The position of these residues not only disrupts the catalytic triad but disrupts the S1 and S1’ sites in a way that prevents the oxyanion hole from forming (Figure 5), the result of which is presumed to result in an inactive form of the protein which is not biologically relevant for our screening efforts. However, monomer A adopting the expected Ser-His-Glu-like catalytic domain structure enables fragment screening to be carried out.

**Figure 5:**
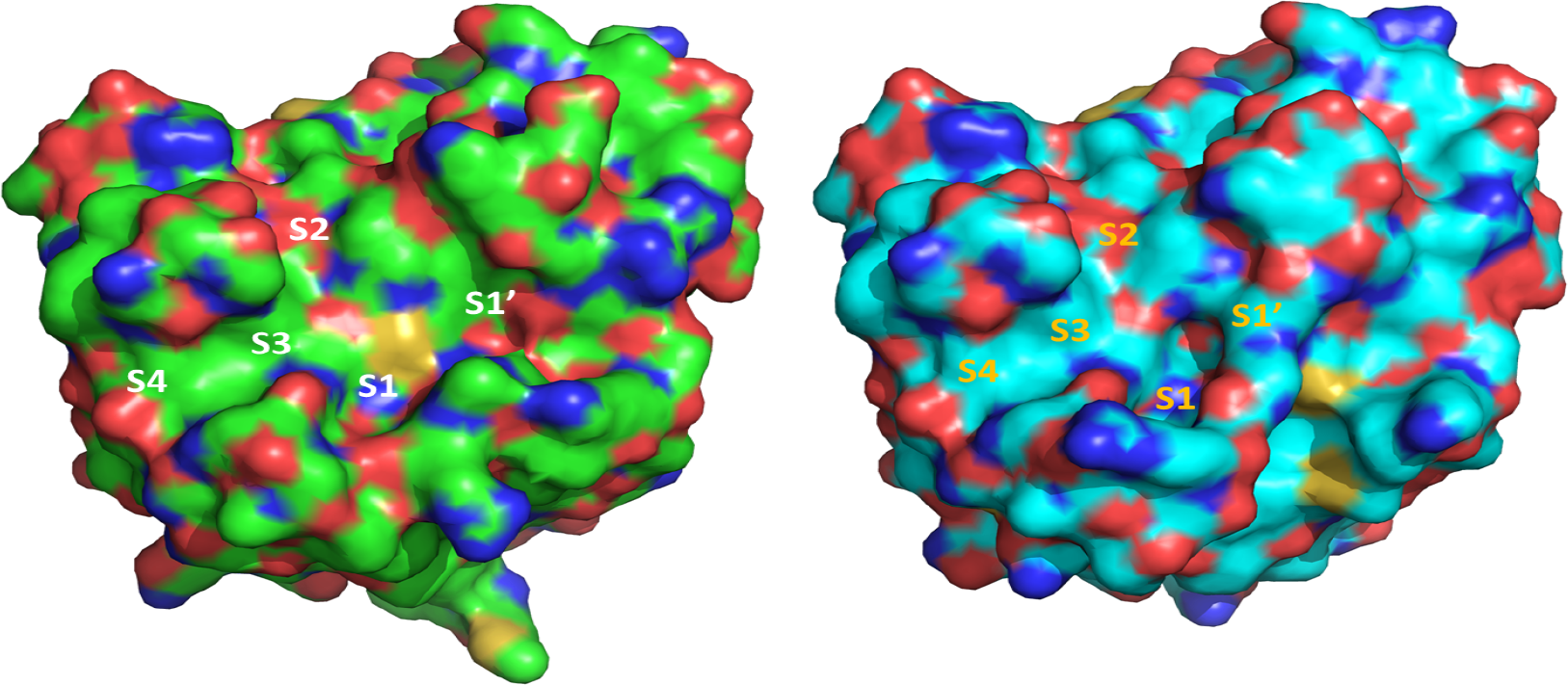
Surface representation of the catalytic sites of monomer A (Green) and B (Cyan). Monomer A has the catalytic triad in the correct conformation and has the expected active site conformation. Monomer B, owing to Cys147 being everted away, adopts a different conformation of the active site where the S1 and S1’ sites are distorted with the Cys147 no longer present and the oxyanion hole no longer present.

### Crystallographic Fragment screen of 3C^pro^

The active site of 3C^pro^ is not involved in any crystal contacts or occluded by crystal packing, making it an ideal system for fragment screening. The crystals were tested for solvent tolerance to DMSO, with the crystals displaying no change in diffraction quality after incubation with 12.5% v/v DMSO for 3 hours allowing the screening of 1,231 fragments from a range of available libraries.

Structural data obtained from this screen is deposited in the PDB but is also disseminated through the Fragalysis web server. Fragalysis set about addressing a major gap in the structure-enabled drug discovery field, namely the lack of a readily accessible informatics tool that allows non-specialist users to reliably progress their fragment hits to on-scale potency. We have therefore developed Fragalysis Cloud (http://fragalysis.diamond.ac.uk), which publicly hosts all XChem’s collaborative fragment results, alongside those of a few friendly users. Its scope is deliberately fragment-specific, encompassing analysis, progression, collaboration, and dissemination. Important features include, pre-curation of fragment structures (so that fragment binding can be directly analysed and compared), shareable views (snapshots) of the data that precisely reconstitute what the user sees and thus streamlines collaboration and compute-heavy algorithms that can be simply executed, and results reviewed and shared, by pre-configured GUI tools. Primary and secondary data is disseminated in Fragalysis following FAIR mechanisms, in downloadable zip archives that include self-documenting PDFs and snapshot URLs to explain the context and history of the download. Data can also be kept private, for collaboration, or made public, for dissemination.

In total, 101 unique 3C^pro^ structures were identified by the PanDDA algorithms as having ligand binding events within the (see Figure 6). Binding of these fragments was observed across both monomers, with the majority of the fragments boundin three main areas; the monomer-monomer interface, a pocket on monomer B around residues Arg19 and Tyr48, and finally the catalytic triad of monomer A.

**Figure 6:**
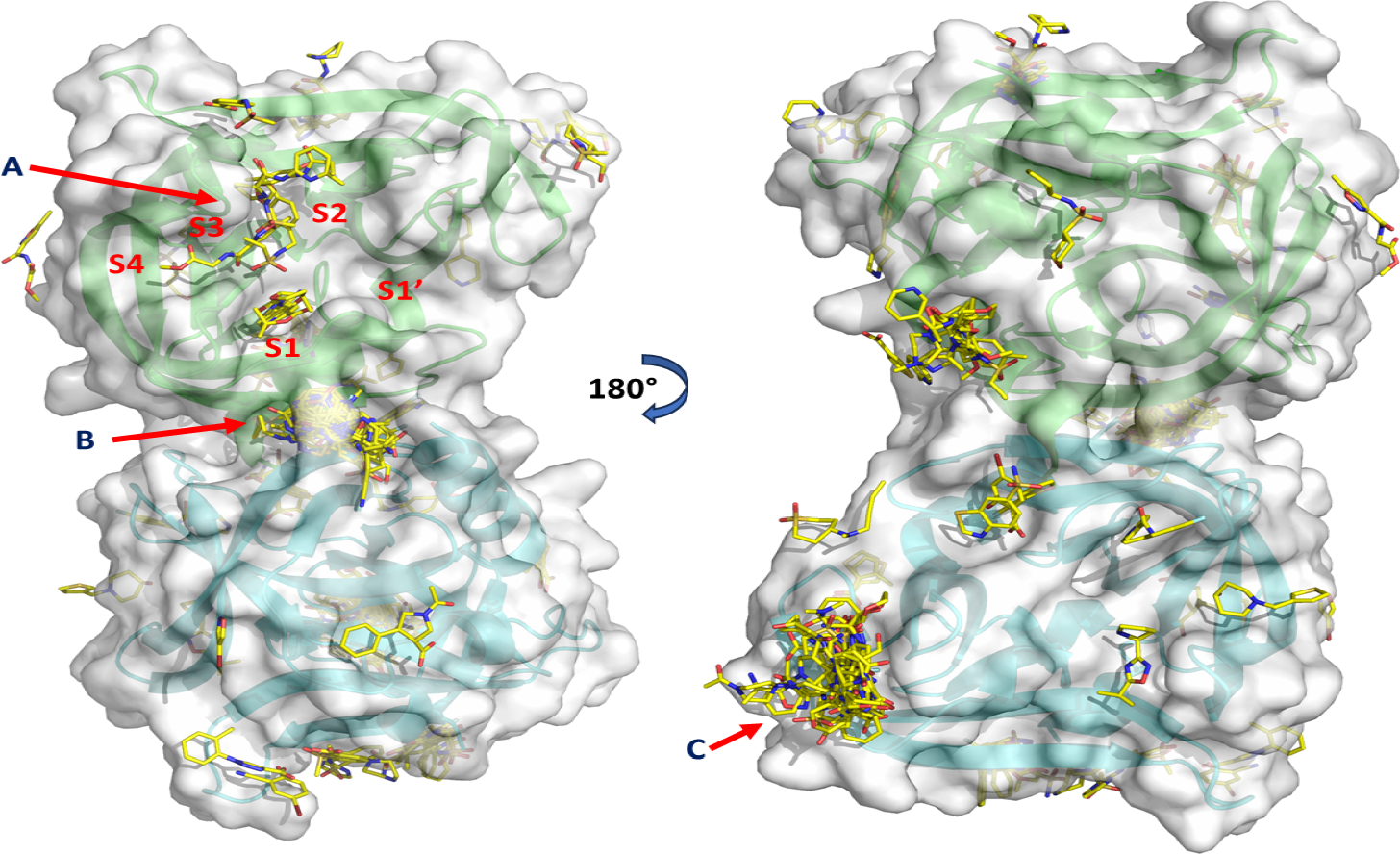
Overview of the 3C^pro^ binding sites on both monomers A (Green) & B (Cyan) for the various bound fragments. The 3 main binding regions are highlighted; **(A)** Monomer A catalytic site, **(B)** Monomer-monomer interface and **(C)** Monomer B Arg19 & Tyr48 pocket.

In total 28 fragments were observed to bind around the monomer A/B interface (Figure 6). This region of the protein is a solvent exposed pocket that contains the N-termini of monomer B and the opposite side of the Gly-X-Cys-Gly-Gly motif which makes up the monomer A catalytic site, containing the catalytic Cys147. The pocket contains the N-terminal α-helix of monomer B, with residues Gly1, Pro2 and Phe3 forming interactions with the fragments either through their side chains or the peptide backbone. Other residues from monomer A also form interactions with the fragments, the main residues being Phe109, Phe140 and Arg143. The arrangement of the 3 aromatic phenylalanine rings within close proximity of each other enables strong π-stacking to occur [56], with many of the fragments that bound here containing an aromatic ring. However, due to the pocket only forming based on the packing of the two monomers, it is likely artefactual and the fragments that bound here are unlikely to be viable for further development.

Another pocket region that is shown in the crystallographic structure of 3C^pro^ occurs around Arg19 and Tyr48 of monomer B, however this pocket is not observed in monomer A (Figure 6). Within the pocket 44 fragments were observed to bind, again with the majority of the fragments containing an aromatic ring. The pocket is composed of several β-sheet loops, with the peptide backbone forming many of the interactions with the fragments. However, there are 2 main residues within this pocket, Arg19 and Tyr48, both of which actively form hydrogen bonds to many of the fragments, as well as suspected π-interactions between the aromatic sections of the fragments and Tyr48. There are no fragments observed in the same site on monomer A even though there are minimal structural differences between the two, other than a slight shift in the positioning of Glu21 and the presence of 2 rotameric states of Arg19. The total lack of fragments is attributed to the crystal packing, whereby a loop from the symmetry mate is occluding the pocket in monomer A, preventing ligand binding.. The presence of the pocket and its relative proximity to the active site leads to speculation that this region may be a potential allosteric site, however deeper analysis into this hypothesis is needed.

The final, and most interesting, binding site identified was the active site on monomer A (Figure 6), which had 12 fragments observed binding across the S1, S2 and S3 subsites(Figure 7).

**Figure 7:**
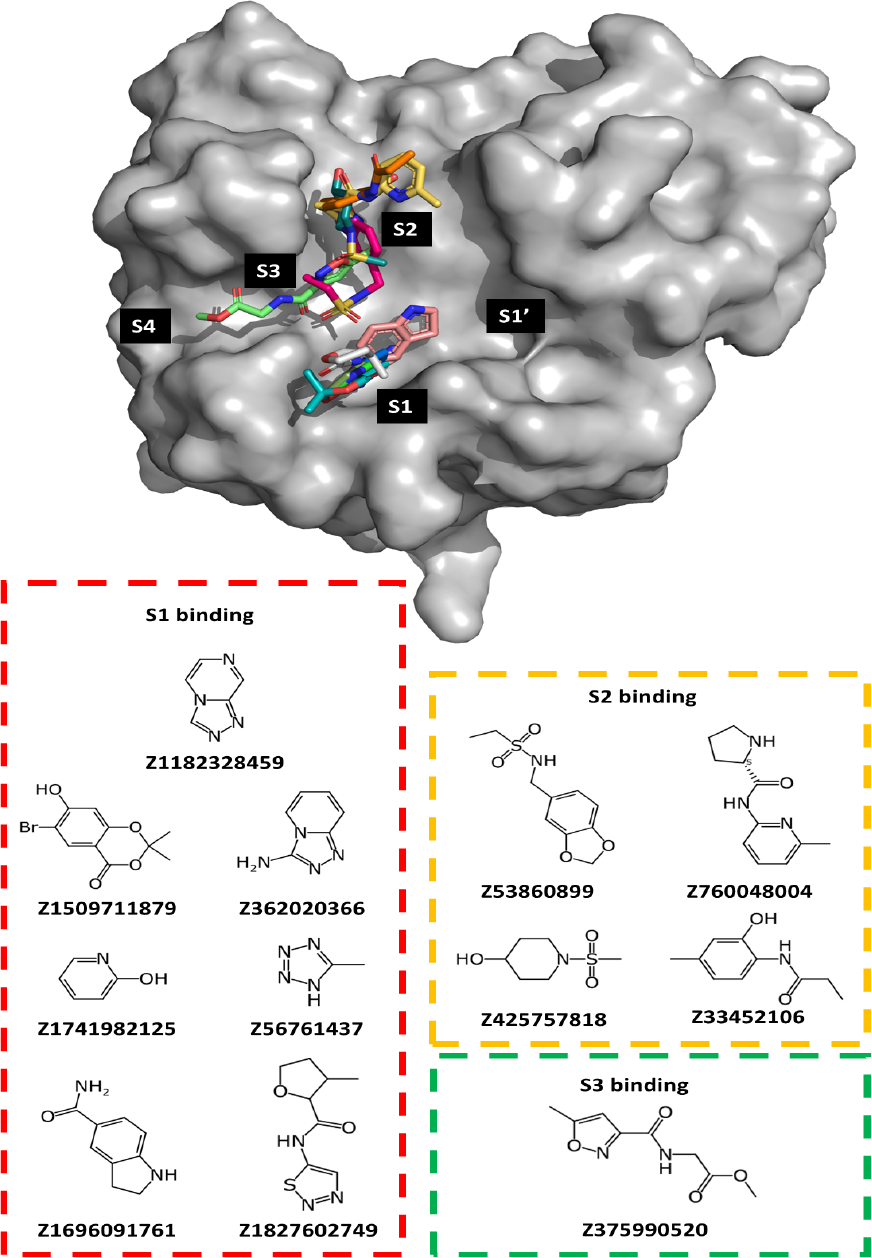
*S*urface structure of 3C^pro^ monomer A showing the position of the 12 observed binding fragments in the various subunits and the chemical structures of the fragments that bound in the S1 (**red Box)**, S2 **(yellow Box)** and S3 **(green box)** subsites.

The fragments that bound in the S1 pocket mimic the terminal amide of the conserved glutamine within the natural substrates. These fragments interact heavily with His161 and Thr142, through the formation of hydrogen bonds, as well as the backbone of a hairpin loop via a bridging water molecule. None of the fragments were found to displace the heavily coordinated water in the Thr142 pocket.

The S1 binding fragments fall into two groups, a pyrazole-like scaffold with a non-protonated sp^2^ nitrogen or an oxygen substituent. The former were a 1,2,3-thiadiazole (Z1827602749), 1,2,4-triazolo[4,3-a]pyrazine (Z1182328459), 1,2-diazaindolizine (Z362020366), and tetrazole (Z56761437). None of these are able to hydrogen bond with the crystalline water. In the latter group one hit, Z1696091761, was a terminal amide akin to the native substrate, Z1741982125 is a pyridone, while Z1509711879 is a hydroxy-benzodioxinone. In the DSi-poised library, there are 57 terminal amides and 2 gamma-lactams with an unsubstituted nitrogen (Z1270255928 and Z1537361757). The library also contained three benzimidazolones (Z26794305, Z26794338 and Z26794351), which did not bind, presumably because the six-membered arene would be unable to be bind in similar manner to the triazolopyrazine hits and thus unable to form pi-amide interactions with Gly164. Gamma-lactam is a common bioisostere of a primary amide used in peptidomimetic design due to the entropic benefit of reduced rotational freedom. For this target, the screen revealed that a pyrazole-like moiety may be privileged and provide a good opportunity for strong binder design.

Only three hits formed interactions with the thiol or thiolate group of the catalytic cysteine, namely Z1509711879 (via a halogen bond), Z1696091761 (via a S·π interaction) and Z362020366 (via a salt bridge), and these hits also featured hydrogen bonding with His161 Figure 8 below shows the S1 bound fragments and the residues involved in interacting with Cys147 and His161.

**Figure 8:**
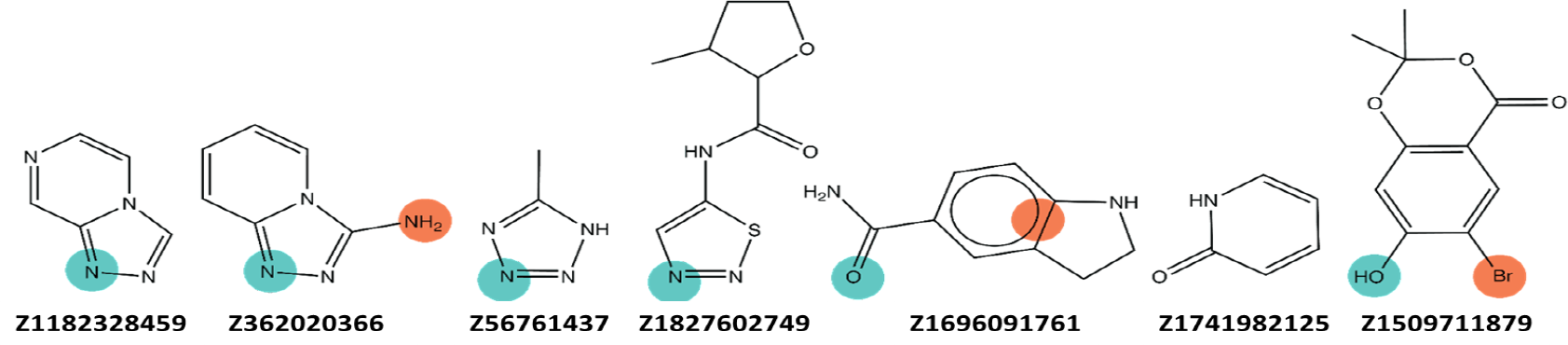
Chemical structures of the fragment hits observed in the S1 sidechain pocket. Hydrogen acceptor with His161 in teal, interacting groups with Cys167 in coral.

The P2 position of the substrate is variable, preferably a large hydrophobic residue (Figure 1). This is reflected in the fragments that bound in the S2 region, which were a diverse set of large compounds containing hydrophobic regions, with the position of His40 leading to minor π-interactions. The backbone interaction of substrate (P3-P2 residues) with the side of the beta-sheet residues Gly163 and Gly164 were mimicked by some hits, in particular sulfonamide (Z53860899). Due to a turn in the substrate sequence these hits are close to hits in the P1 sidechain pocket within the S1 subunit. In light of this, and the absence of P1 backbone pocket hits, a focus of initial fragment elaboration was to bridge the chemical matter in the S1 pocket with that along the S2–S3 backbone, and possibly the S2 pocket.

### Fragment elaboration

The first iteration of fragment elaborations were catalogue-based in order to rapidly and cost-effectively explore diversity via common scaffolds. Initial fragment elaborations were performed via a panel of computational methodologies, either via direct catalogue enumeration or via *de novo* design followed by catalogue analogue search. As detailed below, three direct catalogue enumeration methods were used, namely FragmentKnitwork (substructure search), Arthiorian quest (ambiguous SMARTS pattern search), and OpenEye ROCS (shape-and-colour search). Two additional *de novo* methods were used, Fragmenstein (stitching together of parent hit atoms) and STRIFE (EGNN based generator), which were followed by analogue-by-catalogue search. The crucial principle underpinning these approaches is the preservation of the position of the parent hit fragment as this is a strong predictor for crystallographic success [17], this is especially important as the fragments are typically too weak for meaningful biophysical affinity data to be generated at this point [42].

Whereas common in method benchmarking, ranking by predicted binding potential is a poor metric as it does not capture the diversity of interactions and does not factor in the divergence of the compound to the parent hit —a source of novelty but also risk. Where applicable, the enumerated compounds were not ranked by predicted energy or ligand efficiency, but via a multifactor metric taking into account deviation from the parent hits (RMSD, number of differing atoms and the number of interactions maintained and lost). These were then clustered by interaction, specifically, k-means clustered by residue-level interaction presence scaled by frequency of that interaction in the whole dataset in order to downweigh against clusters characterised by the lack of highly conserved interactions, which is a negative.

The focus on binding conformation of the parent compound, diversity, and on ranking to improve fragment exploration led to 268 derivative compounds designed and ordered. After trialling the curation and selection of algorithmic follow-up compounds by medicinal chemists for real-world drug discovery using Fragalysis, further refinements were made to improve the searchability of uploaded design sets and navigation of the designed compounds for review (**Fig.9**).All compounds were screened against 3C^pro^ using high-throughput crystal soaking with binding of 10 observed with high confidence.

Four derivative compounds preserved the hydrogen-bond acceptor interaction with His161 of their parent hit (Z362020366), via a 1,2-diazaindolizine moiety (Z4466932329, Z3561686360 and Z3484591695) or a 1,2,3-trizole moiety (Z2600009240), emphasising the likely strong enthalpic contribution of this group (**Figure 9**). Z2600009240 is a trizole and an indole bound by a flexible linker, the indole was erroneously predicted to bind the S2 pocket, whereas in the crystal structure its location is an artefact of crystallisation. However, this hit may aid in fragment progression because unlike a 1,2-diazaindolizine, which has limited applicable expansions in catalogue space, a triazole is the reaction product of a Huisgen cycloaddition thus it could allow easy exploration of chemical space to either side of the ring. Another hit that acts as hydrogen-bond acceptor with His161 is the gamma-lactam Z2590396248, which surprisingly bound not with the nitrogen towards the coordinated water as seen with the primary amide or certain lactam containing fragment hits, but away, further indicating that this interaction is not critical despite its presence with the native substrate.

**Figure 9:**
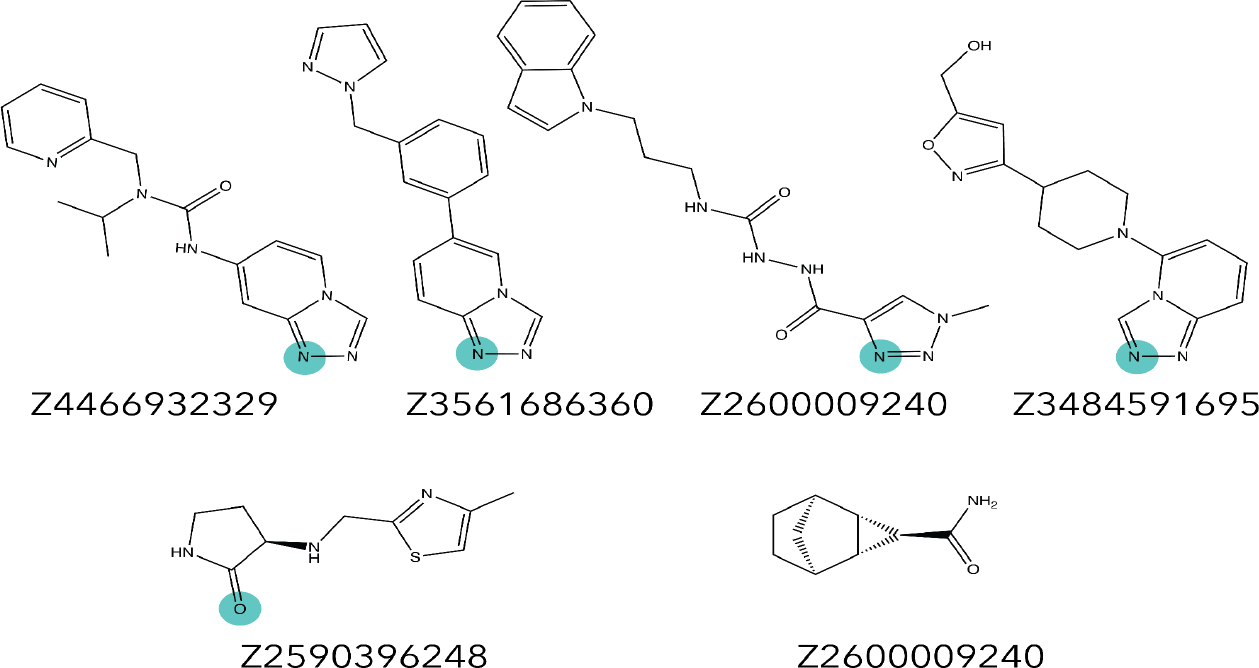
Chemical structure of fragment elaboration hits that were confidently seen in the crystallographic structure. Highlighted is the hydrogen acceptor with His161 in Teal.

The S1 pocket was further populated by a norbornane containing hit enumerated by shape-and-colour (OpenEye ROCS),PV-002320089721, which bound above His161, hinting towards the possibility that some 3D compounds may also be able to bind in the pocket.

It should be noted that several compounds were shortlisted, including Z2590396248 and Z3561686360, as predicted expansions towards the catalytic Cys147, but all were not observed in the crystal structures, possibly due to the mobility of the cysteine. It can be conjectured that this region is flexible in order to allow induced fit of the highly flexible substrate, which would otherwise have a very low binding rate were the catalytic environment rigid (conformational selection), and when the peptide is bound prior to catalysis the strong enthalpic interactions outweigh the entropic cost of rigidification. However, in a fragment screen, as fragment binding is generally assumed to be enthalpically driven due to their size, this would be disadvantageous. Nevertheless, if correct it would suggest that inhibiting the initial substrate binding by occupying the critical sites, such as the His161 interactions, would allow the design of high potency drug-like molecules.

### Future work

It is unsurprising that the region accommodating many of the fragments is the one that accommodates the sidechain of the universally conserved S1 residue (glutamine) of the native substrates. This pocket has a strong preference for triazolo-bicyclic compounds due to hydrogen bonding with His161 and Thr142, presumably mimicking the native ligand’s amide-oxygen head, however, the interaction of the native ligand’s amide nitrogen is not frequently seen. Consequently, further exploration ought to focus on triazolo-bicyclic compounds, or in lieu of these uncommon scaffolds in catalogue space, triazole compounds as a placeholder until the lead optimization stages of the drug discovery campaign where an appropriate bioisostere could be identified.

Expansions towards the catalytic cysteine, Cys147, was unsuccessful, potentially due to the mobility of the region. Consequently, future strategies should focus on linking the S1 and S2 sites not by following the backbone of the native ligand, but by crossing over to S3, potentially elaborating the Z53860899 sulfonamide. This has the added benefit that several facile reactions such as sulfo-Schotten–Baumann reaction would allow nano-scale combinatorial exploration of this space.

Additionally, as elaborations increase in size and presumably improve in affinity, orthogonal biophysical and biochemical assays can be implemented to complement the structural data available.

## Conclusion

This study has focused on the highly conserved enterovirus 3C^pro^. By rapidly performing a crystallographic fragment screen and initial hit-to-lead development we hope to promote confidence that the design of pan-enteroviral inhibitors is possible.

The fragment screen has afforded a set of 102 discrete fragments covering four main regions of the protein. The 12 fragments within the active site engage in interactions with key substrate binding residues, and span the S1, S2 and S3 subsites involved in natural substrate binding. The initial active site fragment hits have been used to design a number of follow-up compounds that elaborate from the parent fragments, with the majority focusing on the fragments that interact with His161 creating a core structure. It is highly likely that further fragment-merging and elaboration can be utilised to efficiently develop compounds that span the entirety of the active site whilst keeping true to the parent hit structures. These future compounds would hopefully achieve high binding potencies and would suitably inhibit the natural function of the protein *in vivo*.

The rapid (3 month) turnaround of the crystallographic fragment screen and open data dissemination (with data being uploaded to both the PDB and Fragalysis), enables wide-spread use of this valuable dataset. Furthermore, the quick turnaround of fragment elaborations (designed and curated within a 9 week timeframe) highlights the efficiency of the ASAP consortium pipeline, and can be applied to other future pre-pandemic targets to ensure equitable pandemic preparedness [2] .

## Supporting information

## Acknowledgements

The authors would like to acknowledge Diamond Light Source for access to the fragment screening facility XChem, for usage of DSi-Poised and other libraries and for beamtime on beamline I04-1 under the proposal LB32627.

We thank Stephanie Wills, Lucy Vost and Matteo Ferla (University of Oxford) for their invaluable contributions to running the various fragment elaboration algorithms and their assistance in the manuscript preparation.

## Author contributions

RML wrote the manuscript. RML & CWET carried out fragment screening and data analysis. DF, JCA, BHB, and PGM supported data processing, data collection and validation. MW, WT and TZT assisted with computational analysis and chemistry. SW, EPW, MF, XN, CG and LK carried out protein expression, purification and quality control. DF, LK and ASG reviewed the manuscript.

## Data and materials availability

All data needed to evaluate the conclusions in the paper are present in the paper and/or the Supplementary Materials. Fragalysis Crystallographic coordinates and structure factors for all structures have been deposited in the PDB with the following accessing codes:

## PDB Codes

8CNX, 7GNV, 7GNW, 7GNX, 7GNY, 7GNZ, 7GO0, 7GO1, 7GO2, 7GO3, 7GO4, 7GO5, 7GO6, 7GO7, 7GO8, 7GO9, 7GOA, 7GOB, 7GOC, 7GOD, 7GOE, 7GOF, 7GOG, 7GOH, 7GOI, 7GOJ, 7GOK, 7GOL, 7GOM, 7GON, 7GOO, 7GOP, 7GOQ, 7GOR, 7GOS, 7GOT, 7GOU, 7GOV, 7GOW, 7GOX, 7GOY, 7GOZ, 7GP0, 7GP1, 7GP2, 7GP3, 7GP4, 7GP5, 7GP6, 7GP7, 7GP8, 7GP9, 7GPA, 7GPC, 7GPD, 7GPE, 7GPF, 7GPG, 7GPH, 7GPI, 7GPJ, 7GPK, 7GPL, 7GPM, 7GPN, 7GPO, 7GPP, 7GPQ, 7GPR, 7GPS, 7GPT, 7GPU, 7GPV, 7GPW, 7GPX, 7GPY, 7GPZ, 7GQ0, 7GQ1, 7GQ2, 7GQ3, 7GQ4, 7GQ5, 7GQ6, 7GQ7, 7GQ8, 7GQ9, 7GQA, 7GQB, 7GQC, 7GQD, 7GQE, 7GQF, 7GQG, 7GQH, 7GQI, 7GQJ, 7GQK, 7GQL, 7GQM, 7GQN, 7GQO, 7GQP, 7GQQ, 7GQR

